# Chimera alert! The threat of chimeric sequences causing inaccurate taxonomic classification of phytoplasma strains

**DOI:** 10.1101/2023.04.10.535501

**Authors:** Bhavesh Tiwarekar, Kiran Kirdat, Shivaji Sathe, Xavier Foissac, Amit Yadav

## Abstract

The 16S rRNA gene is a primary marker used to study bacterial taxonomy, particularly for bacteria with ‘*Candidatus*’ status, where it serves as a primary reference sequence for identification. Despite its usefulness, ensuring the authenticity of publicly available sequences has always posed a challenge. To address this issue, multiple tools and databases with curated collections of reference 16S rRNA gene sequences were established. However, even with these efforts, many sequence chimeras still exist. These chimeric gene sequences are formed by the fusion of sequence parts from multiple organisms and can arise from errors during PCR amplification or sequencing. The objective of this study was to identify chimeric sequences from a dataset of over 4500 phytoplasma 16S rRNA gene sequences using multiple tools. The tools ChimeraSlayer and UCHIME identified 12 and 11 chimeric sequences, respectively. Notably, these sequences included the reference 16S rRNA gene sequence of ‘*Ca*. Phytoplasma wodyetiae’ strain Bangi-2 (KC844879) and ‘*Ca*. Phytoplasma allocasuarinae’ (AY135523). The study’s outcomes indicated the existence of chimeric 16S rRNA gene sequences, emphasizing the threat posed by such sequences in correctly assigning taxonomic status to phytoplasma strains. These findings underscore the importance of rigorously verifying the authenticity of 16S rRNA gene sequences to ensure their accuracy in identifying and classifying bacterial species.

## Introduction

The 16S rRNA gene is a widely used phylogenetic marker for bacterial taxonomy (Woese, 1987), particularly for identifying bacteria with ‘*Candidatus*’ status (Murray and Stackebrandt, 1995; Murray et al., 2020). The term ‘*Candidatus*’ refers to a provisional taxonomic status for bacteria that have not yet been cultured in the laboratory, but their presence has been inferred through molecular markers and techniques used to probe them (Murray and Stackebrandt, 1995). As a result, the 16S rRNA gene became a critical tool for identifying and characterizing ‘not-yet-cultivated’ bacterial taxa, like ‘*Ca*. Phytoplasma’ (IRPCM, 2004; Wei and Zhao, 2022; Kirdat et al., 2023). However, using the 16S rRNA gene as a marker for bacterial taxonomy can be complicated by the presence of chimeric sequences. Chimeric sequences are the result of PCR amplification that merges two or more parent sequences. They can arise when partially extended strands act as primers in subsequent PCR cycles. In the case of phytoplasma chimeric sequences, their formation may be attributed to the presence of other endophytes or the co-infection of the host by two different phytoplasma strains. Chimeric sequences can cause problems in the identification and classification of microbial taxa, especially bacteria. Chimeric sequences are commonly observed in amplicons generated from environmental samples that contain a mixture of DNA from different microorganisms (Wang and Wang, 1996; Smyth et al., 2010; Edgar et al., 2011; Haas et al., 2011; Fonseca et al., 2012).

The growing number of publicly available 16S rRNA gene sequences led to the development of multiple databases, including the Ribosomal Database Project (RDP) (Cole et al., 2007), Greengenes (DeSantis et al., 2006), SILVA database (Pruesse et al., 2007), and EzBioCloud (Yoon et al., 2017), aimed at creating a reliable and authentic database for assigning accurate taxonomic status. These databases contain a vast collection of curated 16S sequences. However, despite extensive curation efforts, many chimeric sequences still exist (Ibal et al., 2019; Johnson et al., 2019; Sze and Schloss, 2019; Abellan-Schneyder et al., 2021). To address this issue, especially chimeric sequences generated from environmental samples, specialized software tools are used to compare the sequence of the PCR product to a reference database of known sequences and identify any regions of the sequence that are likely to be chimeric (Ashelford et al., 2005, 2006; Smyth et al., 2010; Edgar et al., 2011; Haas et al., 2011).

This study analyzed the 16S rRNA gene sequences of phytoplasma strains submitted to the GenBank/DDBJ/EMBL database, along with the reference sequence of provisional phytoplasma species, to identify anomalous sequences, specifically chimeric sequences. The aim was to emphasize the importance of verifying the authenticity of 16S rRNA gene sequences for accurate identification and classification of bacterial species, as the presence of chimeric sequences can impact the assignment of valid taxonomic status to phytoplasma strains

## Methods

The phytoplasma 16S rRNA gene sequences that were >1200 bp, including the reference sequences of provisional phytoplasma species, were downloaded from GenBank/DDBJ/EMBL database and merged to create a single input multi-FASTA file. Duplicate sequences were removed from the dataset using DAMBE v7.2.152 (Xia, 2018), and the orientation of the remaining sequences was verified using Orientation Checker v1.0 (Ashelford et al., 2006). The resulting dataset comprised 4,506 sequences, which were used as input for ChimeraSlayer and UCHIME. The same set of sequences was used for Mallard analysis, adding sequence of *E. coli* (strain K12, U00096) as a default parameter, and these were aligned using the online version of MAFFT v7 (https://mafft.cbrc.jp).

This study utilized four specialized software tools viz. ChimeraSlayer (Haas et al., 2011), UCHIME (Edgar et al., 2011), Mallard (Ashelford et al., 2006), and Pintail (Ashelford et al., 2005) to identify chimeric sequences of phytoplasma 16S rRNA genes available in the GenBank/DDBJ/EMBL database. In addition, the study performed a phylogenetic analysis to confirm the chimeric nature of the identified sequences.

The first tool, ChimeraSlayer, identifies potential chimeras by performing a sequence alignment between query 16S rRNA sequence and a database of reference chimera-free 16S sequences. The top two reference sequences are then selected based on alignment scores. Finally, an evolutionary framework is used to identify a query sequence as a chimera when such a query sequence exhibits greater sequence homology with an in-silico chimera that would form using the selected top reference sequences. The algorithm calculates a score for each putative chimeric sequence based on how likely it is to be a chimeric sequence. The sequence is classified as chimeric if the score exceeds a threshold (Haas et al., 2011).

The UCHIME algorithm generates four non-overlapping segments from the query sequence. Each segment is further used to identify the best matching candidate from the reference chimera-free database. The best two candidates are used in a three-way alignment with the main query sequence. A score is computed based on the entire alignment, and the query sequence is flagged as a chimera based on a pre-determined threshold score (Edgar et al., 2011).

Mallard applies the Pintail algorithm in a many (query) -on-one (subject) comparison. The output of Mallard is a graph where the resultant D.E. (Deviation from Expectation) values are plotted against their corresponding mean observed percentage differences. The Pintail graphs associated with each D.E. value can be further inferred to discern the nature of sequence anomaly. The chimeric sequences are identified based on breakpoints, which are regions of the contig where the sequence switches from one reference sequence to another. The outliers can be identified based on the cut-off line defined by the user and the calculated D.E. values (Ashelford et al., 2006).

The Pintail algorithm aligns the query sequence and a subject sequence. Pintail then evaluates the evolutionary divergence between the two sequences through a sliding window analysis with a pre-specified window size and fixed step size. The resulting deviation from expectation (D.E.) value represents the probability of a chimeric sequence. Pintail generated graphs illustrate the precise locations of chimeric breakpoints and their corresponding reference sequences that were instrumental in identifying them (Ashelford et al., 2005).

The potential chimeric phytoplasma 16S rRNA gene sequences were analyzed phylogenetically using the breakpoint information observed from the Pintail graphs to confirm the results obtained using the chimera detection tools. For this, the reference 16S rRNA gene sequences of phytoplasma provisional species were aligned using ClustalW (Thompson et al., 1994), and two phylogenetic trees were constructed. The phylogenetic trees were constructed using MEGA 7(Kumar et al., 2016).

## Results and Discussion

ChimeraSlayer identified 12 chimeric 16S rRNA gene sequences out of 4,506 sequences, which included two reference sequences ‘*Candidatus* (*Ca*.) Phytoplasma (P.) allocasuarinae’ (AY135523) (Marcone et al., 2004) and ‘*Ca*. P. wodyetiae’ Bangi-2 (KC844879) (Naderali et al., 2017). The chimeras were identified using a minimum bootstrap value of 90%, indicating high confidence in the classification. Moreover, ChimeraSlayer could also determine the parent (left parent A or right parent B) sequences used to create the chimera (Table 1). Additionally, the UCHIME program identified 11 phytoplasma 16S rRNA gene sequences as chimeric, including the two reference sequences mentioned earlier. The chimeras were identified using an empirically determined threshold (h) score obtained from a 3-way alignment of a query sequence with a two-parent sequence. The default ‘h’ score used in the program was 0.3 (Edgar et al., 2011) (Table 2).

**Table 1:** ChimeraSlayer output representing identified potential chimeras in 4,506 sequences used for the analysis and their respective parents. The minimum bootstrap value used for calling a sequence chimeric is 90 %. ^1^ PerIDQLAQRB: Similarity of the query to the left side of parent A and right side of parent B; ^2^BootStrapA: confidence in the query as a chimera related to the left side of parent A, right side of parent B; ^3^PerIDQLAQRB: similarity of the query to the right side of parent A and left side of parent B; ^4^BootStrapB: confidence in the query as a chimera related to the right side of parent A and left side of parent B.

| Phytoplasma strain | Parent A | Parent B | PerIDQL<br>AQRB<sup>1</sup> | Boot<br>StrapA<sup>2</sup> | PerIDQL<br>BQRA<sup>3</sup> | Boot<br>StrapB<sup>4</sup> |
| --- | --- | --- | --- | --- | --- | --- |
| 'Wodyetia bifurcata' yellow decline<br>phytoplasma clone 1 (KY069029) | 'Ca. Phytoplasma asteris' AYWB<br>(M30790) | Ca. Phytoplasma cynodontis' LW01<br>(AJ550984) | 90.43 | 0 | 99.20 | 100 |
| Sugarcane phytoplasma strain<br>Mauritius (AJ539180) | 'Ca. Phytoplasma asteris' AYWB<br>(M30790) | 'Ca. Phytoplasma dypsis' (MT536195) | 97.02 | 96.3 | 90.50 | 0 |
| 'Ca. Phytoplasma **wodyetiae**'<br>(KC844879) | 'Ca. Phytoplasma asteris' AYWB<br>(M30790) | 'Ca. Phytoplasma cynodontis' LW01<br>(AJ550984) | 88.04 | 0 | 96.23 | 99.8 |
| 'Ca' Phytoplasma solani' isolate<br>Lorestan (KF130968) | 'Ca. Phytoplasma australasia'<br>PpYC (Y10097) | 'Ca. Phytoplasma solani' 2642BN<br>(AJ964960) | 89.66 | 0 | 98.80 | 100 |
| 'Phoenix dactylifera' phytoplasma<br>isolate Jor2 (MT880242) | 'Ca. Phytoplasma australasia'<br>PpYC (Y10097) | 'Ca. Phytoplasma ulmi' EY1<br>(AY197655) | 96.86 | 99.8 | 89.77 | 0 |
| 'Phoenix dactylifera' phytoplasma<br>isolate Jor8 (MT880244) | 'Ca. Phytoplasma australasia'<br>PpYC (Y10097) | 'Ca. Phytoplasma ulmi' EY1<br>(AY197655) | 95.80 | 99.5 | 89.26 | 0 |
| 'Phoenix dactylifera' phytoplasma<br>isolate Jor7 (MT880243) | 'Ca. Phytoplasma australasia'<br>PpYC (Y10097) | 'Ca. Phytoplasma ulmi' EY1<br>(AY197655) | 97.33 | 100 | 89.83 | 0 |
| 'Phoenix dactylifera' phytoplasma<br>isolate Jor9 (MT880245) | 'Ca. Phytoplasma australasia'<br>PpYC (Y10097) | 'Ca. Phytoplasma ulmi' EY1<br>(AY197655) | 95.63 | 99.7 | 88.34 | 0 |
| 'Phoenix dactylifera' phytoplasma<br>clone Jor5 (MT956596) | 'Ca. Phytoplasma australasia'<br>PpYC (Y10097) | 'Ca. Phytoplasma ulmi' EY1<br>(AY197655) | 96.19 | 100 | 88.41 | 0 |
| 'Phoenix dactylifera' phytoplasma<br>clone Jor5 (MT956597) | 'Ca. Phytoplasma australasia'<br>PpYC (Y10097) | 'Ca. Phytoplasma ulmi' EY1<br>(AY197655) | 95.46 | 99.9 | 87.34 | 0 |
| 'Phoenix dactylifera' phytoplasma<br>isolate SAHail2 (MT956950) | 'Ca. Phytoplasma australasia'<br>PpYC (Y10097) | 'Ca. Phytoplasma ulmi' EY1<br>(AY197655) | 95.30 | 99.6 | 88.74 | 0 |
| 'Ca. Phytoplasma **allocasuarinae**'<br>(AY135523) | 'Ca. Phytoplasma australasia'<br>PpYC (Y10097) | 'Ca. Phytoplasma pyri' PD1<br>(AJ542543) | 89.98 | 0 | 97.38 | 99.6 |

<sup>1</sup>PerIDQLAQRB: Similarity of the query to the left side of parent A and right side of parent B; <sup>2</sup>BootStrapA: confidence in the query as a chimera related to left side of parent A and right side of parent B; <sup>3</sup>PerIDQLAQRB: similarity of the query to the right side of parent A and left side of parent B; <sup>4</sup>BootStrapB: confidence in the query as a chimera related to right side of parent A and left side of parent B.

**Table 2:** UCHIME output showing identified potential chimeras and their respective parents used for the analysis of 4,506 phytoplasma 16S rRNA gene sequences of length >1200 bp. The default threshold score of 0.3 was used to determine chimeras. Higher scores indicate a stronger chimeric signal. ^1^Score: Score calculated to report chimera (Default value: 0.3); ^2^IdQM: Percent identity of the query (Q) and the model (M) constructed as a segment of Parent A and a segment of Parent B; ^3^IdQA: Percent identity of Q and A; ^4^IdQB: Percent identity of Q and B; ^5^IdAB: Percent identity of A and B.

| Phytoplasma strain | Parent A | Parent B | Score<sup>1</sup> | IdQM<sup>2</sup> | IdQA<sup>3</sup> | IdQB<sup>4</sup> | IdAB<sup>5</sup> |
| --- | --- | --- | --- | --- | --- | --- | --- |
| 'Wodyetia bifurcata' yellow decline phytoplasma clone 1 (KY069029) | 'Ca. Phytoplasma cynodontis' LW01 (LT558776) | 'Ca. Phytoplasma asteris' AYWB (M30790) | 4.45 | 99.2 | 96.5 | 93.1 | 90.8 |
| 'Ca. Phytoplasma **wodyetiae**' (KC844879) | 'Ca. Phytoplasma cynodontis' LW01 (LT558776) | 'Ca. Phytoplasma asteris' AYWB (M30790) | 4.13 | 99.1 | 95.9 | 93.7 | 90.9 |
| 'Ca. Phytoplasma solani' strain Lorestan (KF130968) | 'Ca. Phytoplasma solani' 2642BN (AJ964960) | 'Ca. Phytoplasma australasia' PpYC (Y10097) | 1.92 | 98.8 | 97.1 | 91.8 | 89.1 |
| 'Phoenix dactylifera' phytoplasma isolate SAHail2 (MT956950) | 'Ca. Phytoplasma australasia' PpYC (Y10097) | 'Ca. Phytoplasma ulmi' EY1 (AY197655) | 1.65 | 98.1 | 91.2 | 96.3 | 90 |
| Brinjal little leaf phytoplasma, Clone TN-43 (KP027525) | *Escherichia coli* K12 (U00096) | 'Ca. Phytoplasma trifolii' C.P. (AY390261) | 1.53 | 85.3 | 78.1 | 79.3 | 71.4 |
| 'Ca. Phytoplasma **allocasuarinae**' (AY135523) | 'Ca. Phytoplasma mali' AT (CU469464) | 'Ca. Phytoplasma australasia' PpYC (Y10097) | 1.43 | 97.4 | 95 | 92.5 | 90.7 |
| Sugarcane phytoplasma 16S rRNA gene, strain Mauritius (AJ539180) | 'Ca. Phytoplasma asteris' AYWB (M30790) | 'Ca. Phytoplasma dypsidis' RID7692 (MT536195) | 1.05 | 97.7 | 94.1 | 94.5 | 91.1 |
| 'Prunus persica' phytoplasma Meerut (HM988985) | 'Ca. Phytoplasma asteris' AYWB (M30790) | 'Ca. Phytoplasma trifolii' C.P. (AY390261) | 0.70 | 96.9 | 94.6 | 94 | 91.3 |
| 'Phoenix dactylifera' phytoplasma isolate Jor2 (MT880242) | 'Ca. Phytoplasma australasia' PpYC (Y10097) | 'Ca. Phytoplasma ulmi' EY1 (AY197655) | 0.69 | 98.1 | 92.6 | 96.9 | 90.3 |
| Brinjal little leaf phytoplasma, Clone TN-45 (KP027526) | *Escherichia coli* K 12 (U00096) | 'Ca. Phytoplasma trifolii' C.P. (AY390261) | 0.52 | 75.5 | 71 | 73.1 | 69.2 |
| 'Phoenix dactylifera' phytoplasma isolate Jor9 (MT880245) | 'Ca. Phytoplasma australasia' PpYC (Y10097) | 'Ca. Phytoplasma ulmi' EY1 (AY197655) | 0.48 | 96.8 | 90.9 | 96 | 90.1 |

The 16S rRNA gene sequence of ‘*Ca*. P. wodyetiae’ Bangi-2 (KC844879) was analyzed for potential chimerism using the Pintail algorithm. The algorithm compared the sequence with its putative parent sequences, ‘*Ca*. P. cynodontis’ LW01 (LT558776) (Marcone et al., 2004; Kirdat et al., 2021) and ‘*Ca*. P. asteris’ AYWB (M30790) (Lee et al., 2004), resulting in two profiles (Figure 1). Profile A compared Bangi-2 with AYWB, while Profile B compared Bangi-2 with LW01. Both profiles showed close matches between the aligned sequences, depicting the parental segments. Next, the Pintail algorithm compared Bangi-2 with each putative parent sequence, resulting in deviation from expectation (D.E.) values of 4.25 with AYWB and 3.51 with LW01. Based on the comparison with the reference strains, the probability of two non-anomalous sequences producing D.E. values of 4.25 and 3.51 and differing by 7.84% and 4.42%, respectively, was estimated to be p<0.001, indicating a strong anomaly in the sequence (Figure 1).

**Figure 1.**
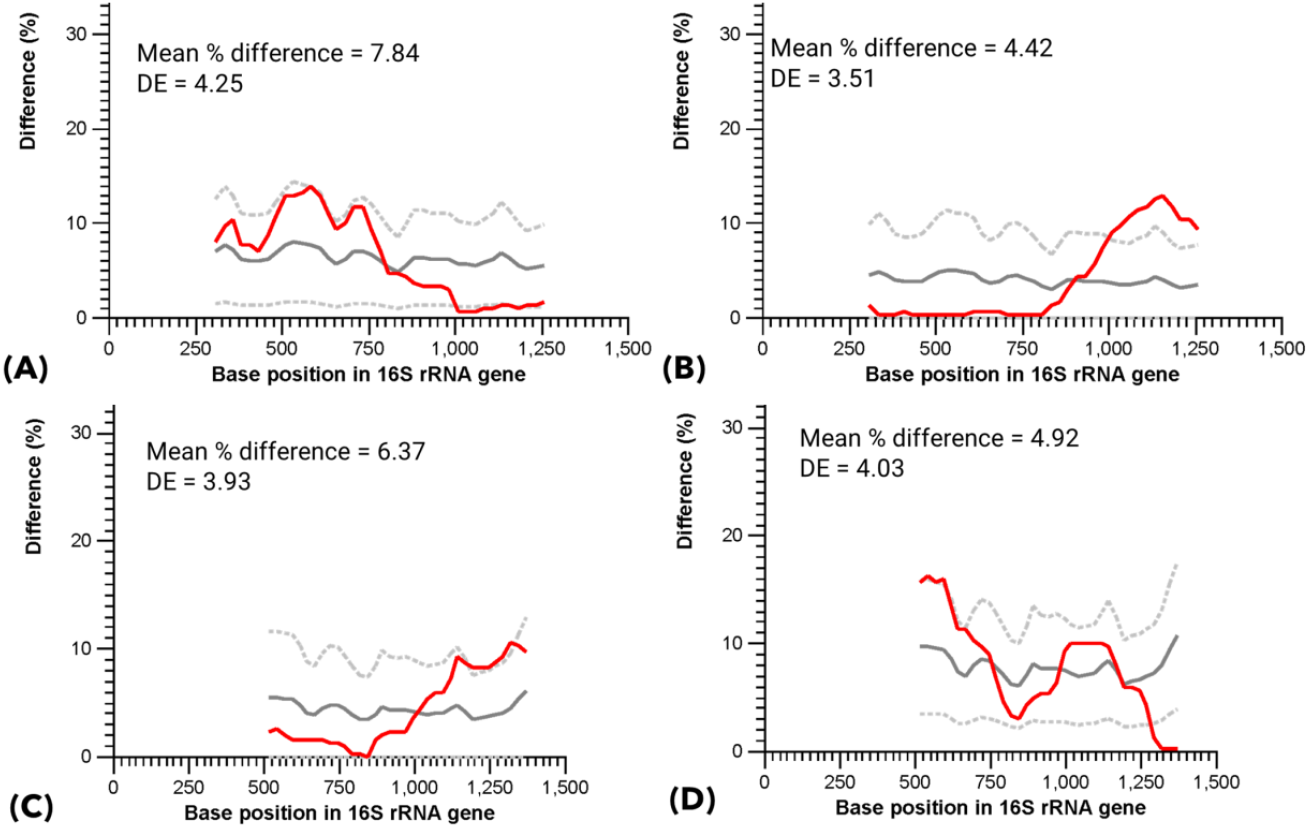
Pintail profile for ‘*Ca*. Phytoplasma wodyetiae’ Bangi-2 (KC844879) compared with ‘*Ca*. Phytoplasma asteris’ AYWB (M30790) (A) and ‘*Ca*. Phytoplasma cynodontis’ LW01 (LT558776) (B); Pintail profile of ‘*Ca*. Phytoplasma allocasuarinae’ (AY135523) compared with ‘*Ca*. Phytoplasma australasia’ PpYC (Y10097) (C) and ‘*Ca*. Phytoplasma rhamni’ BAWB (X76431) (D). The graphs show variation in % difference between aligned sequences, determined with a 300-base window, moving 25 bases simultaneously along the sequences’ length. Solid grey line indicates the expected percent difference; dotted grey lines indicate plus or minus five expected percent differences, and the red line indicates the observed percent difference.

In a similar fashion, the Pintail algorithm was used to analyse the 16S rRNA gene sequence of ‘*Ca*. P. allocasuarinae’ (AY135523) along with its putative parent ‘*Ca*. P. australasiaticum’ [formerly ‘Ca. P. australasia’, PpYC (Y10097)] (White et al., 1998; Oren, 2017), which showed a deviation from expectation (D.E.) value of 3.93, differing by 6.37%. Another putative parent, ‘*Ca*. P. rhamni’ BAWB (X76431) (Marcone et al., 2004) had a D.E. value of 4.03, differing by 4.92% (p<0.001), indicating that the reference 16S rRNA sequence of ‘*Ca*. P. allocasuarinae is likely chimeric (Figure 1C). These findings suggest that the reference sequences are potentially chimeric.

Based on the breakpoint information observed in the Pintail graphs, phylogenetic analysis was conducted to investigate the potential chimeric phytoplasma 16S rRNA gene sequences. Neighbour-Joining (N.J.) phylogenetic sub-trees were inferred from the analysis of reference 16S rRNA gene sequences of published phytoplasma species using the first 797 bp (Figure 2, A) and 807-1250 bp (Figure 2, B) using Mega 7. The phylogenetic tree analyses for ‘*Ca*. P. wodyetiae’ Bangi-2 (KC844879) confirmed its close phylogenetic association with its parent strains ‘*Ca*. P. cynodontis’ LW01 (LT558776) (Figure 2, A), and ‘*Ca*. P. asteris’ AYWB (M30790) (Figure 2, B). The PCR assay performed by Naderali et al. in 2017 (description of ‘*Ca*. P. wodyetiae’) identified three co-infecting phytoplasma strains. This phytoplasma was detected in a foxtail palm (*Wodyetia bifurcata*) exhibiting symptoms of ‘Foxtail Palm Yellow Decline (FPYD) ‘ disease. The palm was infected with two known phytoplasmas, closely related to ‘*Ca*. P. asteris’ and ‘*Ca*. P. cynodontis’. The third strain was proposed as a novel provisional species that is potentially chimeric, as suggested by the analyses conducted here. Similarly, the phylogenetic analysis revealed that ‘*Ca*. P. allocasuarinae’ (AY135523) is closely associated with ‘*Ca*. Phytoplasma rhamni’ BAWB (X76431) for the first segment of the 16S rRNA gene (Figure 3, A), as seen in the breakpoint on the Pintail graph. However, the second half of the sequence showed a distinct phylogenetic relationship with ‘*Ca*. Phytoplasma australasia’ PpYC (Y10097) of the PWB phytoplasma group (Figure 3, B). As both parent strains belong to phylogenetically distinct clades within the phylogenetic tree of all provisional phytoplasma species, the putative chimeric nature of the 16S rRNA sequence (AY135523) is suggested.

**Figure 2.**
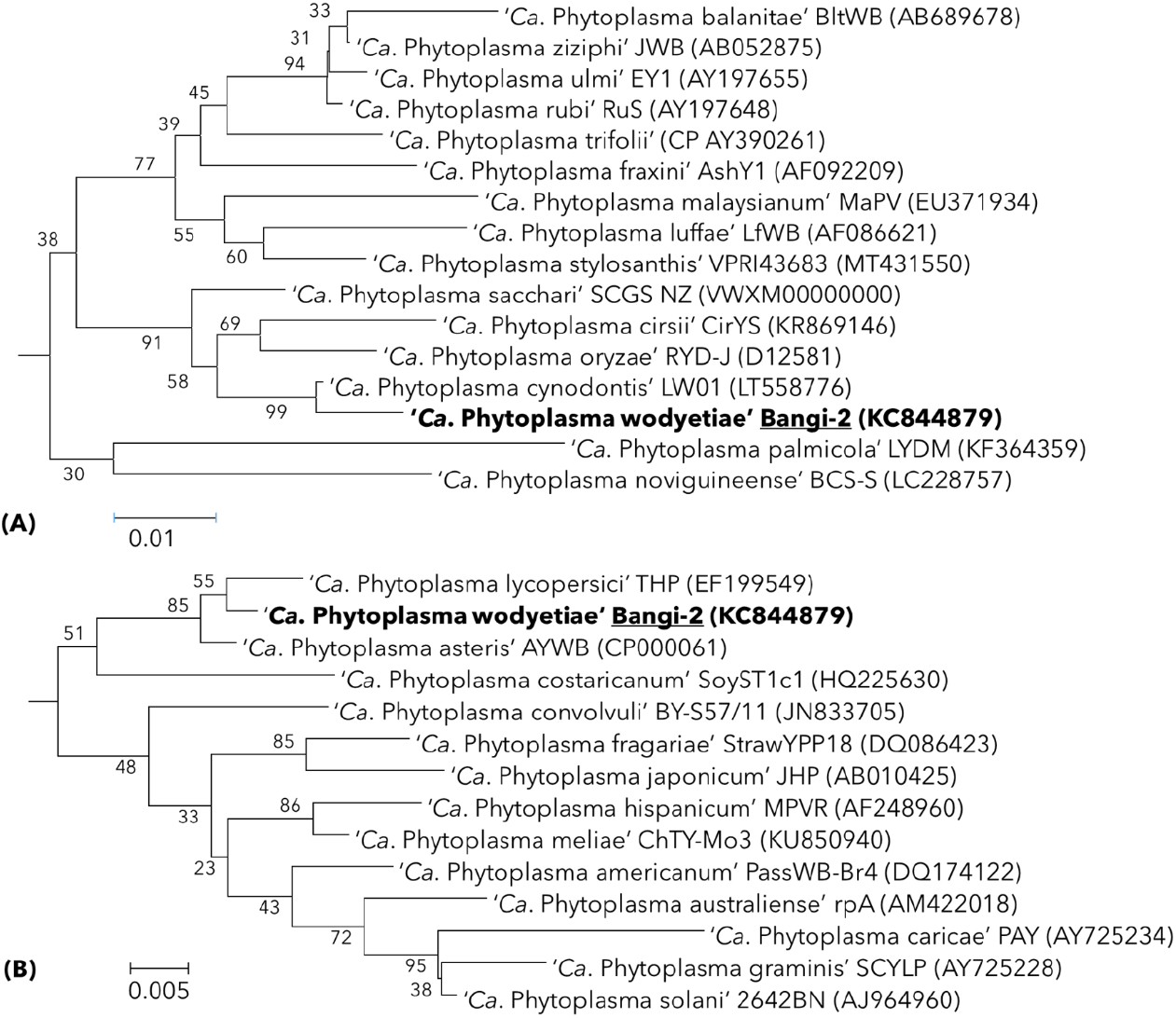
Neighbour-Joining (N.J.) phylogenetic sub-trees inferred from analysis of reference 16S rRNA gene sequences of published phytoplasma species using first 797 bp (A) and 807-1250 bp (B) built using Mega 7 where trees were evaluated by bootstrap analysis based on 1000 replicates. Trees show the phylogenetic position of Bangi-2 (KC844879) in the clade of Aster Yellow (AY) phytoplasmas (A) and Rice Yellow Dwarf (RYD) phytoplasmas (B).

**Figure 3.**
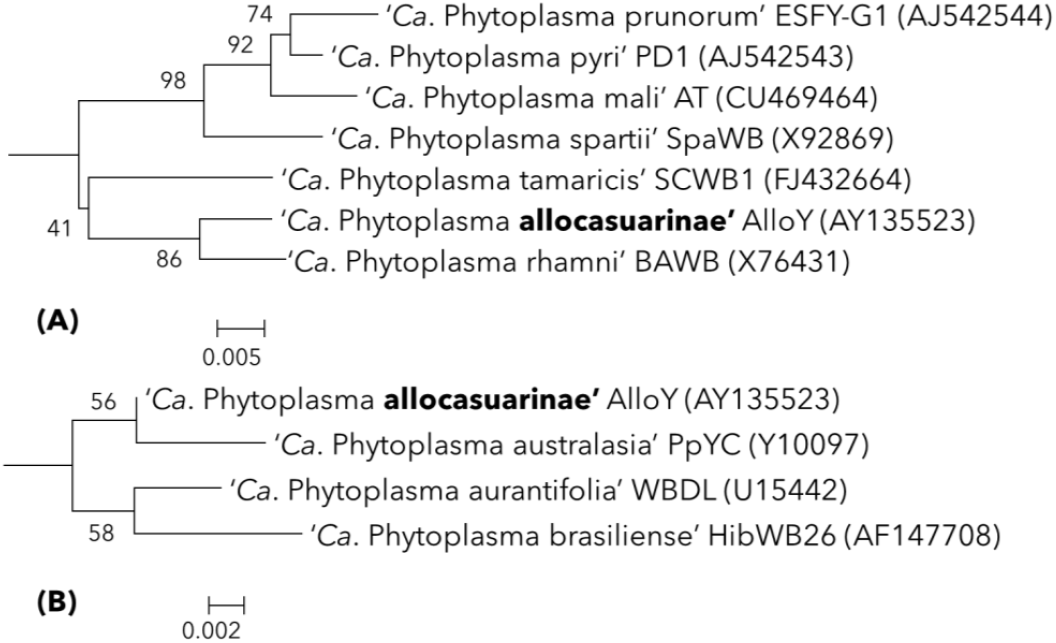
Neighbour-Joining (N.J.) phylogenetic sub-trees inferred from analysis of reference 16S rRNA gene sequences of published phytoplasma species using first 800 bp (A) and 801-1250 bp (B) built using Mega 7 where trees were evaluated by bootstrap analysis based on 1000 replicates. Trees show the phylogenetic position of AlloY (AY135523) in the clade of ‘*Ca*. P. tamaricis’ and ‘*Ca*. P. rhamni’ (A) and Peanut Witches’ broom (PWB) phytoplasmas (B).

In addition to analysing the phylogenetic position of ‘*Ca*. P. allocasuarinae’ and ‘*Ca*. P. wodyetiae’, unusual terminal branching of species ‘*Ca*. P. graminis’ (Pin et al., 2005), ‘*Ca*. P. caricae’ (Pin et al., 2005), and ‘*Ca*. P. lycopersici’ (Arocha et al., 2007) from their ancestor was observed (Figure 4). Although the reference sequences of these strains did not score positive for being chimeric, the multiple sequence alignment revealed the insertion of atypical sequences in their 16S rRNA gene sequence. The probable reasoning behind these insertions could not be determined and may include PCR anomalies. Notably, no other phytoplasma strains related to the five species mentioned above have been reported before or after. These species have only been represented by their 16S rRNA or other gene sequences. This suggests that either these five species may be unique and distinct from other known phytoplasmas or have originated from poor sequencing quality of their 16S rRNA gene (Kirdat et al., 2023).

**Figure 4.**
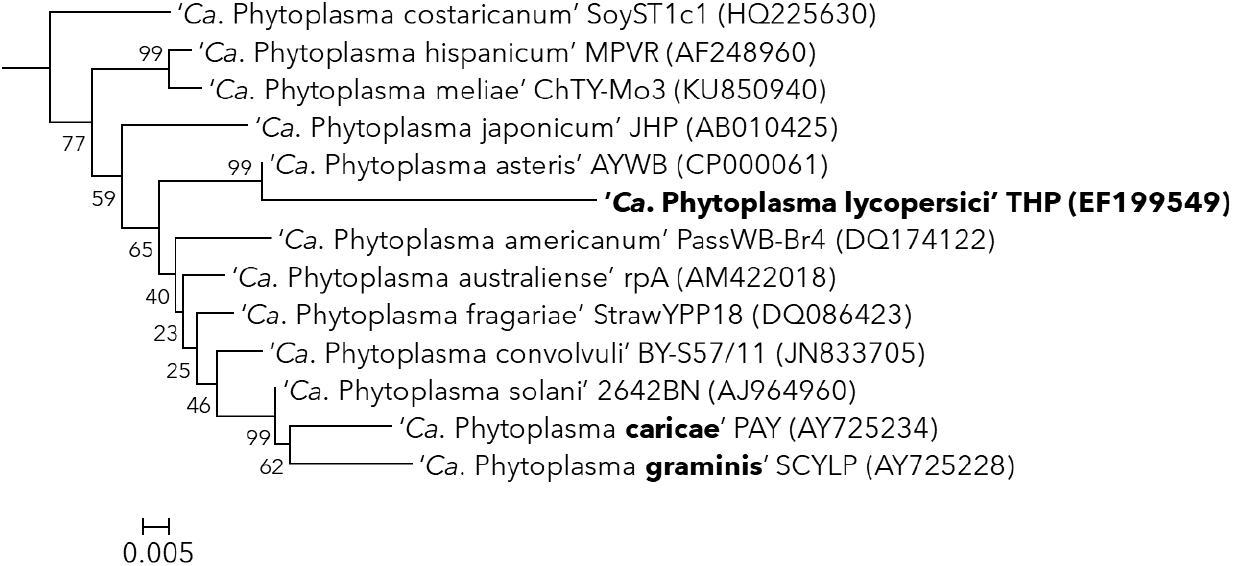
Neighbour-Joining (N.J.) phylogenetic trees inferred from analysis of reference 16S rRNA gene sequences (1250 bp) of published phytoplasma species built using Mega 7 where trees were evaluated by bootstrap analysis based on 1000 replicates. The tree shows the unusual branching of THP, PAY, and SCYLP strains.

The authenticity of the 16S rRNA gene sequence is of utmost importance as it will keep serving as the formal representative (reference sequence) of the proposed provisional species of ‘*Ca*. Phytoplasma’. When characterizing the taxonomy of cultivable bacteria, a polyphasic approach is mandatory, which includes physiological and biochemical characterization (Sneath, 1992). In 2002, the International Committee on Systematics of Prokaryotes (ICSP) further recommended that genome sequence analysis should be included in the polyphasic characterization of novel bacterial species (Stackebrandt et al., 2002). Since then, genome sequence analysis has become a standard component of the polyphasic characterization of novel bacterial species. Therefore, the sequence information from the 16S rRNA gene and percent similarity with existing species should only be considered as a preliminary marker, in addition to the ecological and endemic nature of the phytoplasma strain, to distinguish it at the species level. Holistic consideration should be given to the information gathered from the whole genome, including Overall Genome Relatedness Index (OGRI) values and the endemic nature of its plant and insect host, to propose a new phytoplasma species. Naming and classifying phytoplasma strains to provisional species level can be improved by utilizing comprehensive OGRI values generated from genome sequences of the phytoplasma strains and other features related to their ecological status (Kirdat et al., 2023). The provisional species status should be assigned to strains representing a pool of ecologically evolved populations.

The unavailability of biological material or genome sequences, particularly for provisional species, has led to issues of reproducibility and the inadvertent creation of orphan species (Kirdat et al., 2023). Many phytoplasma strains with provisional species status have been published with limited sequence information, often consisting only of the 16S rRNA gene or a few housekeeping genes. Moreover, a significant proportion of genome sequences deposited in the GenBank database lack supporting Sequence Archive Reads (SRA) data. This situation poses a significant challenge and limits the scope of future taxonomic studies (Wei and Zhao, 2022). Therefore, peer-reviewing the authenticity and validity of sequencing data in scientific research articles plays a critical role in ensuring the reproducibility of the sequencing data. It is imperative to include comprehensive genome sequence information, including SRA data, in taxonomic studies of novel bacterial species to enhance the accuracy and reproducibility of the data. This approach will enable better resolution of taxonomic issues and facilitate more comprehensive comparative analyses among bacterial species.

## Author Approvals

All authors read the manuscript and approved it. The manuscript hasn’t been accepted or published elsewhere.

## Distribution/Reuse Options

Creative Commons Attribution-NonCommercial license (CC-BY-NC): Allowed to distribute, remix, and build upon the work for non-commercial purposes only, as long as appropriate credits are given to the original author(s).

